# Molecular architecture of the ciliary tip revealed by cryo-electron tomography

**DOI:** 10.1101/2023.01.03.522627

**Authors:** T. Legal, M. Tong, C. Black, M. Valente Paterno, J. Gaertig, K.H. Bui

## Abstract

Cilia are essential organelles that protrude from the cell body. Cilia are made of a microtubule-based structure called the axoneme. In most types of cilia, the ciliary tip is distinct from the rest of the cilium. Here, we used cryo-electron tomography and subtomogram averaging to obtain the structure of the ciliary tip of the ciliate *Tetrahymena thermophila*. We show the microtubules in the tip are highly cross-linked with each other and stabilised by luminal proteins, plugs and cap proteins at the plus ends. In the tip region, the central pair lacks the typical projections and twists significantly. By analysing cells lacking a ciliary tip-enriched protein *CEP104/FAP256* by cryo-electron tomography and proteomics, we discovered candidates for the central pair cap complex and explain potential functions of *CEP104/FAP256*. These data provide new insights into the function of the ciliary tip and inform about the mechanisms of ciliary assembly and length regulation.

## Introduction

Cilia are microtubule-based cellular protrusions that extend from the surface of most cells in the human body. Cilia are conventionally classified into two categories: motile cilia and primary cilia. Motile cilia propel cells such as sperm cells or move fluids such as mucus in the respiratory tract, while primary cilia function as sensory organelles (reviewed in (Satir and Christensen 2007).

Due to their important roles, mutations in ciliary proteins can cause multiple disorders called ciliopathies which can lead to infertility, brain malformation or respiratory problems and affect about 1 in 2,000 individuals (Quinlan, Tobin et al. 2008, Reiter and Leroux 2017).

The microtubules inside the cilia assemble into a core structure known as the axoneme. They are templated from the basal body, a circular structure of nine triplet microtubules. The basal body is followed by a transition zone in which the triplet microtubules become doublets made of a complete A-tubule and a partial B-tubule (Ichikawa, Liu et al. 2017). The doublet microtubules become singlet microtubules at the ciliary tip as the B-tubules terminate (Fisch and Dupuis-Williams 2011). Motile cilia also contain two microtubule singlets in the centre, known as the central pair (CP) or central apparatus (Gibbons and Grimstone 1960). The CP microtubules are called C1 and C2 as they differ significantly in structure. C1 is chemically more stable than C2 and the projections on C1 are longer than on C2 (Hopkins 1970). Ciliary beating is driven by outer and inner dynein arms (Lin and Nicastro 2018). They are found on the outside of doublets and slide them apart when activated. The sliding motion is then converted into the bending motion of the cilia because of the nexin-dynein regulatory complex linking the adjacent doublet microtubules (Summers and Gibbons 1971). Doublets are linked to the CP by the radial spokes which coordinate dynein activity (Smith and Sale 1992).

The ciliary tip region has special importance because it is the zone of ciliary growth (Rosenbaum, Moulder et al. 1969). Tubulin and other ciliary subcomplexes are transported to the tips of cilia by intraflagellar transport (IFT) trains where they assemble to build the cilia (Qin, Diener et al. 2004, Craft, Harris et al. 2015). Anterograde trains (walking towards the tip) are reorganised and converted to retrograde trains (walking towards the base) at the ciliary tip (Chien, Shih et al. 2017). In *Chlamydomonas reinhardtii*, the anterograde trains move on the B-tubule while retrograde trains move on the A-tubule of doublet microtubules within the doublet zone of the axoneme (Stepanek and Pigino 2016).

Most of our current knowledge about the ciliary tip comes from early negative-staining experiments. These showed that the CP is capped by a protein complex that looks like a ball attached to the membrane preceded by two plates (Dentler 1984). The spherical plate directly capping the two microtubules is attached to two structures that go inside the microtubules (Suprenant and Dentler 1988). The ends of A-tubules are plugged by a Y-shaped filament bundle that extends 70 nm inside the lumens, namely A-tubule plugs (Suprenant and Dentler 1988). Interestingly, these cap and plug structures are conserved among the unicellular organisms *Chlamydomonas, Tetrahymena* and mammalian tracheal cilia but are not found at the tips of human sperm cells nor human primary cilia (Soares, Carmona et al. 2019). Therefore, the organisation of the ciliary tip may differ depending on the functional specialisations of cilia. The functions of ciliary caps remain to be explored. Also, it is unknown how tubulin subunits are added to the plus ends of microtubules while the capping structures are present. The tips of *Chlamydomonas* cilia were recently investigated using cryo-electron tomography (cryo-ET). Interestingly, the structure of the *Chlamydomonas* tip complex appears very different from that published previously, appearing as a much smaller complex. This study also shows that most microtubules end in a flared conformation near the membrane (Höög, Lacomble et al. 2014). The identities of the proteins of ciliary caps remain to be discovered.

In addition, studies have shown that the CP structure changes in the tip region. In *Tetrahymena*, CP projections present along most of the axoneme terminate at the level where the B-tubules and radial spokes end. In this species, the tip CP region is about 500-nm-long (Sale and Satir 1977). In *Chlamydomonas*, the tip CP region is much shorter, around 200 nm (Pratelli, Corbo et al. 2022). The structures of both doublet-zone CP and the tip region CP in *Tetrahymena* remain to be determined.

Several proteins have been identified to localise to the tips of cilia. Kinesins from kinesin-4, -8 and -13 subfamilies localise to the ciliary tips where they regulate microtubule length (Piao, Luo et al. 2009, Niwa, Nakajima et al. 2012, Wang, Piao et al. 2013, He, Subramanian et al. 2014, Vasudevan, Jiang et al. 2015, Schwarz, Lane et al. 2017). EB1, a microtubule plus-end binding protein, has been shown to localise to the tips of *Chlamydomonas* (Pedersen, Geimer et al. 2003). TOG domain proteins CHE-12/Crescerin, CEP104/FAP256 were also shown to localise to the ciliary tips of the motile cilia in *Tetrahymena* (Louka, Vasudevan et al. 2018). CEP104/FAP256 also localises to the tips of primary cilia in humans (Satish Tammana, Tammana et al. 2013). TOG domains bind to tubulin and promote microtubule elongation (Al-Bassam, Larsen et al. 2007, Das, Dickinson et al. 2015). Interestingly, CEP104/FAP256 is conserved in mammalian primary cilia where it promotes ciliary elongation (Yamazoe, Nagai et al. 2020). Mutations in CEP104/FAP256 cause Joubert syndrome, a brain malformation (Srour, Hamdan et al. 2015). In *Tetrahymena*, knock out of *FAP256* reduces cell swimming velocity and ciliary beat frequency (Louka, Vasudevan et al. 2018). How all these proteins come together to regulate ciliary length and the ends of A- and B-tubules is still unclear.

In recent years, multiple structures of cilia were published. Using single-particle cryo-electron microscopy (cryo-EM), the structures of the doublet microtubules, the radial spokes, the outer dynein arms and the CP were solved to high resolution (Ichikawa, Liu et al. 2017, Ichikawa, Khalifa et al. 2019, Ma, Stoyanova et al. 2019, Khalifa, Ichikawa et al. 2020, Grossman-Haham, Coudray et al. 2021, Gui, Farley et al. 2021, Gui, Ma et al. 2021, Kubo, Yang et al. 2021, Rao, Han et al. 2021, Walton, Wu et al. 2021, Gui, Wang et al. 2022, Han, Rao et al. 2022). At slightly lower resolution, cryo-ET has been used to study the structures of doublet and singlet microtubules in mammalian sperm and identify proteins *de novo* (Chen, Shiozaki et al. 2022, Gui, Croft et al. 2022, Leung, Roelofs et al. 2022). Importantly, the tips of cilia have not been studied at high resolution apart from the singlet-A-tubule of sperm cells (Gui, Croft et al. 2022, Leung, Roelofs et al. 2022).

Here, we used cryo-ET and subtomogram averaging to obtain the structure of the ciliary tip of the wild type and *FAP256A/B-knockout* (KO) *Tetrahymena thermophila* mutant. We show the three-dimensional organisation of the intact ciliary tip, elaborating on the cap, plugs and tip microtubule interactions. We also show that the CP is highly-stabilised by multiple proteins binding to the inside and outside of its microtubules. In addition, using MS analysis of *FAP256A/B-KO* mutants, we generated a list of potential candidates for the CP cap complex and describe potential roles of FAP256.

## Results

### The ciliary tip of *Tetrahymena* is structurally different from the rest of the axoneme

To obtain a high-resolution three-dimensional view of the intact ciliary tip, we deciliated *Tetrahymena thermophila* cells and lightly cross-linked the cilia before plunge-freezing and collecting cryo-electron tomograms (Table S1, Fig. S1A). We collected ∼150 tomograms of ciliary tips and ciliary narrowing regions in three datasets, two with the membrane intact for visualisation of membrane connections and one without the membrane for better visualisation of microtubules. Segmentation of these tomograms after subtomogram averaging (see Materials & Methods) allowed us to observe the organisation of the CP, the singlet A-tubules and the CP cap complex within the ciliary tip (Fig. 1A).

**Figure 1.**
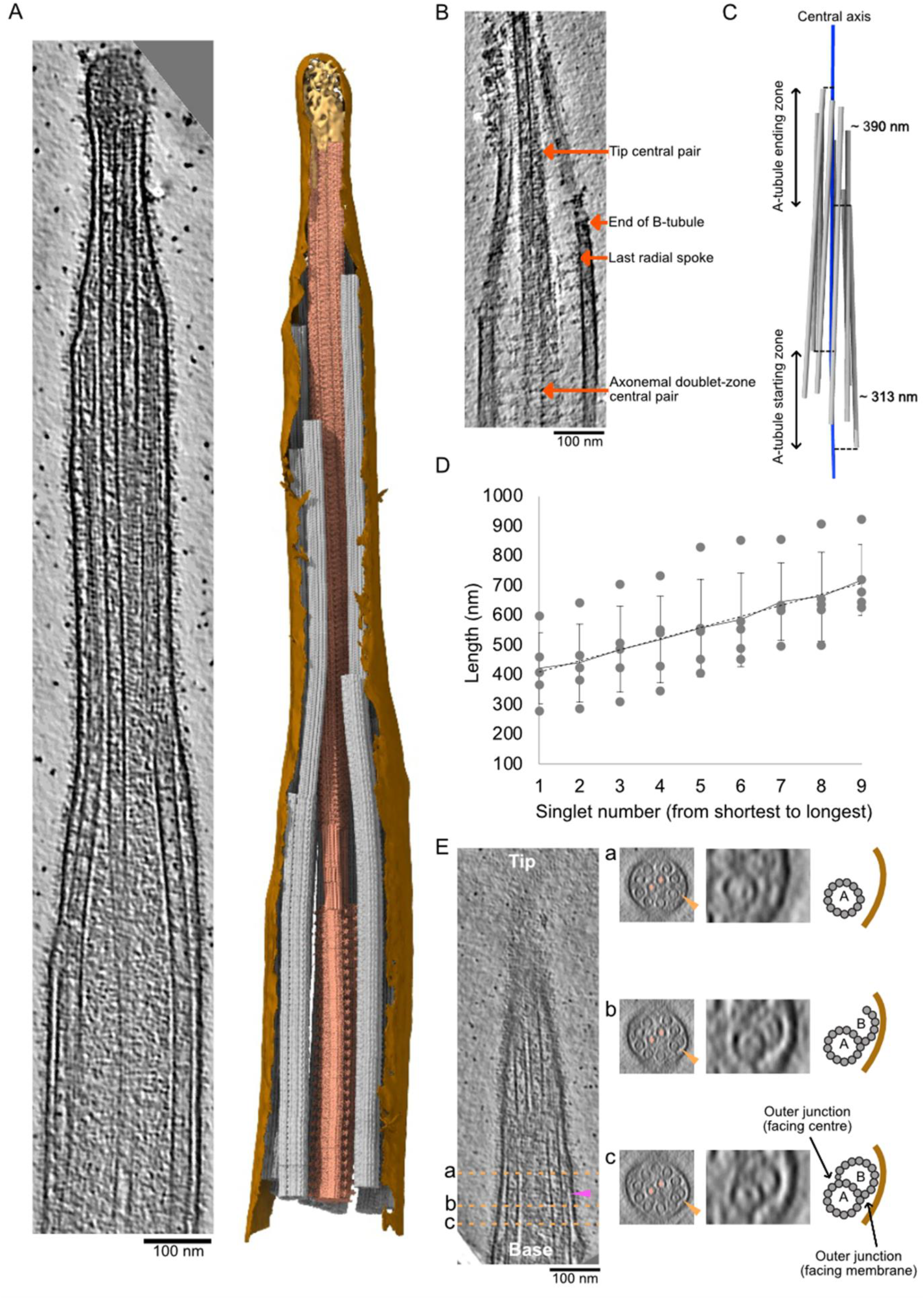
Overview of the ciliary tip. (A) Tomogram slice of a representative ciliary tip. 3D model of the same cilium generated with the subtomogram averages described in this study. Membrane is in brown, the CP is in dark salmon, doublets and singlet A-tubules are in grey, cap complex is in gold. (B) Representative slice of a demembranated cilium showing the tip CP, the end of one B-tubule, the last radial spoke and the axonemal doublet-zone CP. Scale bar: 100 nm. (C) 3D representation of the A-tubules (grey) from a representative tomogram. The central axis of the cilium (centre of the CP) is in blue. (D) Length of A-tubules numbered from shortest to longest. The dotted line shows a linear trend line going through all points. Each dot represents an A-tubule. Error bars show standard deviation. (E) Longitudinal slice through a representative tomogram at the narrowing zone. Orange dotted lines show cross-sections shown in panels a, b and c. Orange arrows point towards the disassembling/incomplete B-tubule. Insets show a doublet with a disassembling/incomplete B-tubule. The CP microtubules are marked by a salmon diamond. Cartoons indicate what is shown in the insets. Purple arrow shows a filament capping the end of the B-tubule.

Denoising these tomograms allowed us to visualise the narrowing zones and ciliary tips in detail (Fig. 1B). We could reveal the structural differences in the CP organisation between the tip region versus the rest of the axoneme. The characteristic projections on the CP stop just before the distance between C1 and C2 microtubules decreases. Within the tip region, the projections of the CP are replaced by a characteristic short spike protein which repeats every 8 nm. Doublet microtubules become singlets when reaching the ciliary tip. The B-tubules, radial spokes and dynein arms all terminate within a short distance from each other (Fig. 1B). The ends of the B-tubules coincide with a narrowing of the cilium diameter, which we refer to as the narrowing zone.

Doublets do not become singlets at the same level. By projecting the start position of each singlet A-tubule (equivalent to the end of each B-tubule) to the cilium’s central axis (Fig. 1C, dotted lines), we measured the A-tubule starting zone to span over 313.3 ± 53.8 nm. Similarly, the A-tubule ending zone showed substantial variation between cilia with an average of 389.7nm ± 272.5 nm. The length of singlet A-tubules also varies significantly from 89 to 720 nm (Fig.1D). Interestingly, when the A-tubules are ordered from shortest to longest, their lengths increase linearly. We did not find any correlation between the A-tubule length and their starting and ending points. Finally, some B-tubules separate from the inner junction (Fig.1E, Fig.S1B). Furthermore, some B-tubules end with a filamentous plug (Fig.1E, purple arrow).

### The tip microtubules interact with many proteins

We next focussed on protein densities specific to the ciliary tip region, which we summarised in a model figure (Fig. 2A). We first examined the ends of A-tubules and CP microtubules. A-tubules ended with slightly curved protofilaments (Fig. 2B, S2A), similar to the plus-ends of microtubules observed *in situ* in axons or fibroblasts (Koning, Zovko et al. 2008, Foster, Ventura Santos et al. 2022). Interestingly, the plug seen at the end of A-tubules appears to be composed of multiple filamentous proteins that extend 110 ± 30 nm inside the A-tubule. We did not observe the “ball” seen in previous negative staining experiments (Suprenant and Dentler 1988). The filaments of the plug make connections with the membrane and the CP microtubules (Fig. 2B, S2A purple arrows). The ends of CP microtubules were difficult to visualise because of the ciliary cap. In most cases, we believe both microtubules are roughly the same length. The CP microtubule cap appears as a mesh of entangled proteins (Fig. 2B, golden circle). We did not observe the plug structures nor the characteristic plates described in previous negative staining studies (Suprenant and Dentler 1988). There are filamentous proteins that extend from the cap to the inside of CP microtubules with an average length of 127 ± 34.7 nm (n=10) (Fig. 2C, S2B).

**Figure 2.**
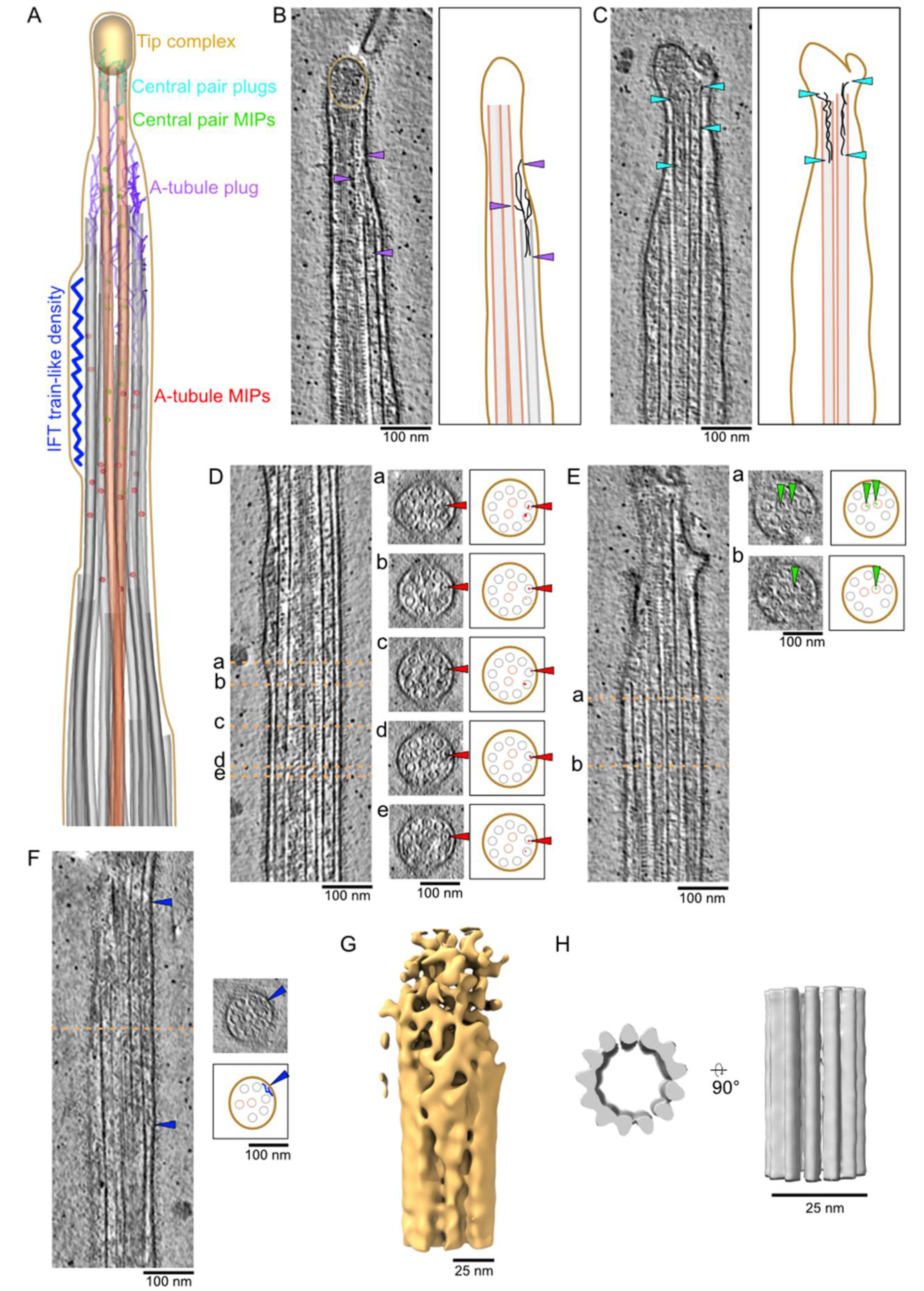
The ciliary tip contains many microtubule-associated proteins. (A) 3D model of the various tip proteins shown in this figure. (B, C) representative tomogram slices along with corresponding diagrams highlighting what is shown in the tomograms. Arrows point to A-tubule plug (B) and CP plug (C). (D, E, F) representative tomogram slices showing A-tubule MIPs (D), CP MIPs (E) and IFT train-like particles (F) along with cross sections indicated by the dotted orange lines. Diagram highlight what is shown in the cross sections. (B - F) Scale bars: 100 nm. (G, H) Subtomogram averages of the cap complex (G) and singlet A-tubule (H). Scale bars: 25 nm.

We also observed a large number of microtubule inner proteins (MIPs) in the A-tubules. Unlike in the A-tubules of the doublet zone, these MIPs do not seem to have a repeating pattern. Some were associated with the microtubule wall while others were inside the microtubule lumen (Fig. 2D, S2C). Many of these MIPs are similar in size and shape to the MIPs observed in mammalian primary cilia (Kiesel, Alvarez Viar et al. 2020) and *Drosophila melanogaster* neurons (Foster, Ventura Santos et al. 2022). In addition, we observed MIPs in both CP microtubules also either associated with the microtubule wall or floating inside the lumen (Fig. 2E, S2D).

IFT trains convert from anterograde to retrograde direction at the ciliary tip (Chien, Shih et al. 2017). IFT train-like particles were bound to the A-tubule in five tomograms (Fig. 2F, S2E). While IFT trains on the A-tubules were found in primary cilia (Kiesel, Alvarez Viar et al. 2020), this is the first observation of IFT trains on the A-tubules in motile cilia. Due to the insufficient number of particles in our tomograms, we could not carry out subtomogram averaging on IFT trains.

To gain insights into the structure of the CP cap, we carried out subtomogram averaging. The resulting average has a resolution of around 100 Å and reveals that the CP cap is asymmetrical (Fig. 2G, table S2).

It has been reported that A-tubules in primary cilia are decorated by a protein that repeats every 8 nm, interpreted by the authors as the microtubule tip-binding protein EB1 (Kiesel, Alvarez Viar et al. 2020). We did not observe any similar decorations on the outside of A-tubules in our tomograms (Fig. 2D, S2C). To confirm this observation, we used subtomogram averaging to obtain a structure of the A-tubule. Our average structure did not reveal MIPs nor MAPs fully decorating the outside of the A-tubules in the tip at this resolution (Fig. 2H, Table S2).

While carrying out subtomogram averaging of the A-tubules, we observed densities on the outside of some A-tubules (Fig. 3A-E). As these densities are not found on all A-tubules and are positioned differently on each A-tubule, they were not seen in the final subtomogram average of all A-tubules. We observed two types of densities: proteins that seem to link two A-tubules (Fig. 3B and C, S3A-C) and proteins that link the A-tubule to the membrane (Fig. 3D and E, S3D). Due to the limited number of particles, the periodicity of these proteins is difficult to determine but some repeat every 8 nm (Fig. 3B, C, E, arrows). Interestingly, we only observed these densities along short portions of the A-tubules. These observations led us to ask whether the A-tubules in the tip were maintained parallel to each other by these links. To answer this question, we measured the distance between all adjacent A-tubules. We found that the average distance between two A-tubules is 31 nm and is constant along the length of the A-tubules (Fig. 3F). Therefore, we believe that these linker densities act as spacers that maintain the distance between the A-tubules in the tip.

**Figure 3.**
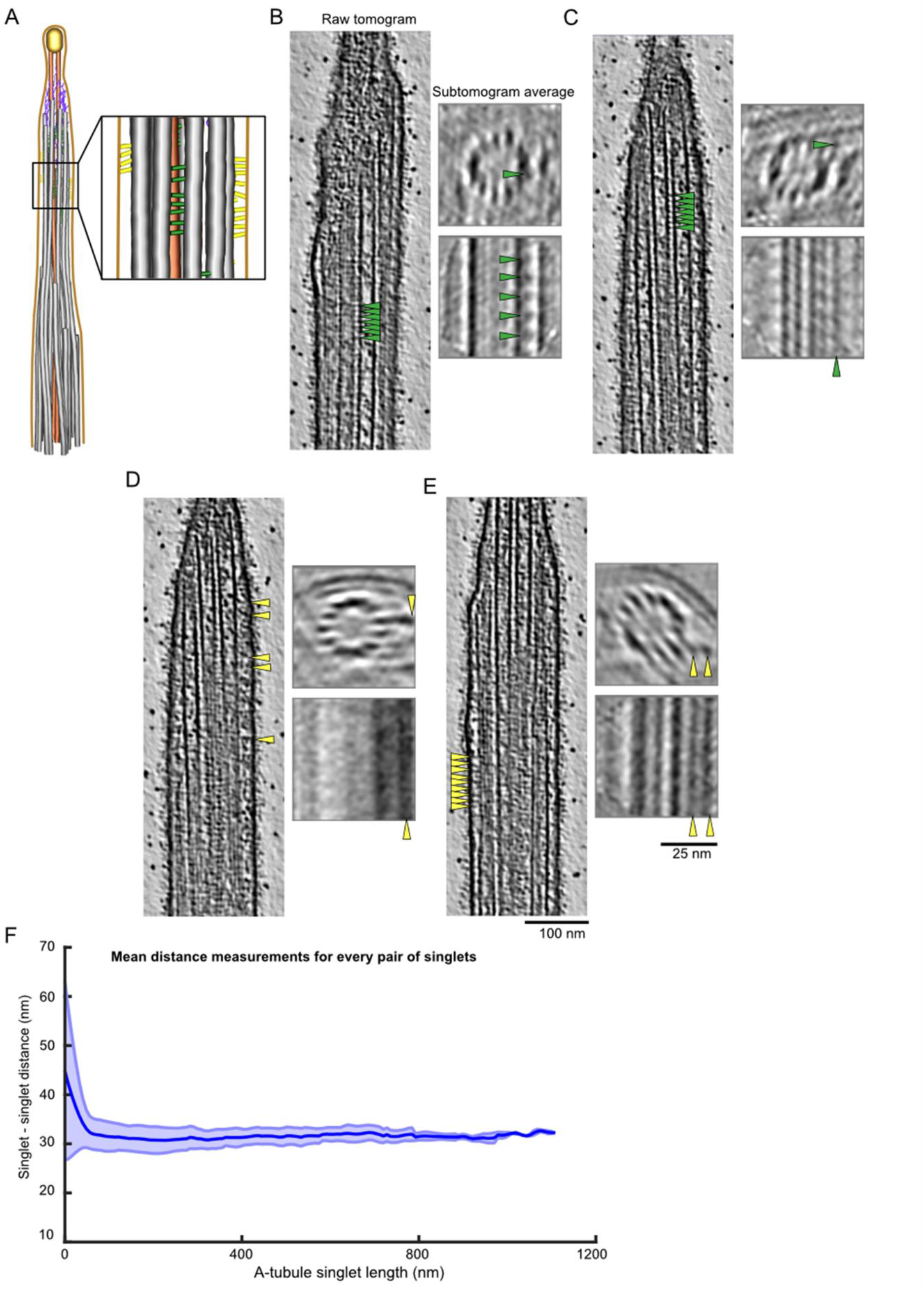
The A-tubules are linked to the membrane and to each other in the ciliary tip. (A) 3D model of the ciliary tip with inset showing A-tubule links in green and membrane links in yellow. (B-E) representative raw tomogram slices along with cross (top) and longitudinal (bottom) slices of subtomogram averages of an individual A -tubule. Green arrows point to A-tubule links (B, C) and yellow arrow point to membrane links (D, E). Scale bars tomograms: 100 nm, subtomogram averages: 25 nm. (F) Graph showing the distance between pairs of A-tubule singlets in the ciliary tip (n = 45).

### The tip CP microtubules are strongly linked to each other

The difference in CP morphology observed in our tomograms (Fig. 1A and B) before and after the narrowing zone motivated us to obtain subtomogram averages of both the doublet-zone CP (hereafter referred to as “main CP”) and the tip CP (Fig. 4A and B, Fig. S4A, Table S2). We obtained a subtomogram average of the 32-nm repeat of the main CP at a resolution of around 23 Å (Fig. 4A, S4B). It shows the canonical architecture of known CP structures (Carbajal-González, Heuser et al. 2013, Fu, Zhao et al. 2019, Leung, Roelofs et al. 2021). The CP of *Tetrahymena thermophila* is similar to the mammalian CP in that the C1f projection is missing. When comparing our structure to other species, we found that the *Tetrahymena* CP is most similar to sea urchin CP (Carbajal-González, Heuser et al. 2013), especially when comparing C1d and C1c projections (Fig. S4C). C2 has similar projections to the known CP structures. C1 and C2 are made up of 13 protofilaments each and are linked by a bridge. *Tetrahymena* CP does not contain big globular MIPs inside C1 nor C2 as opposed to pig and horse CPs (Leung, Roelofs et al. 2021).

**Figure 4.**
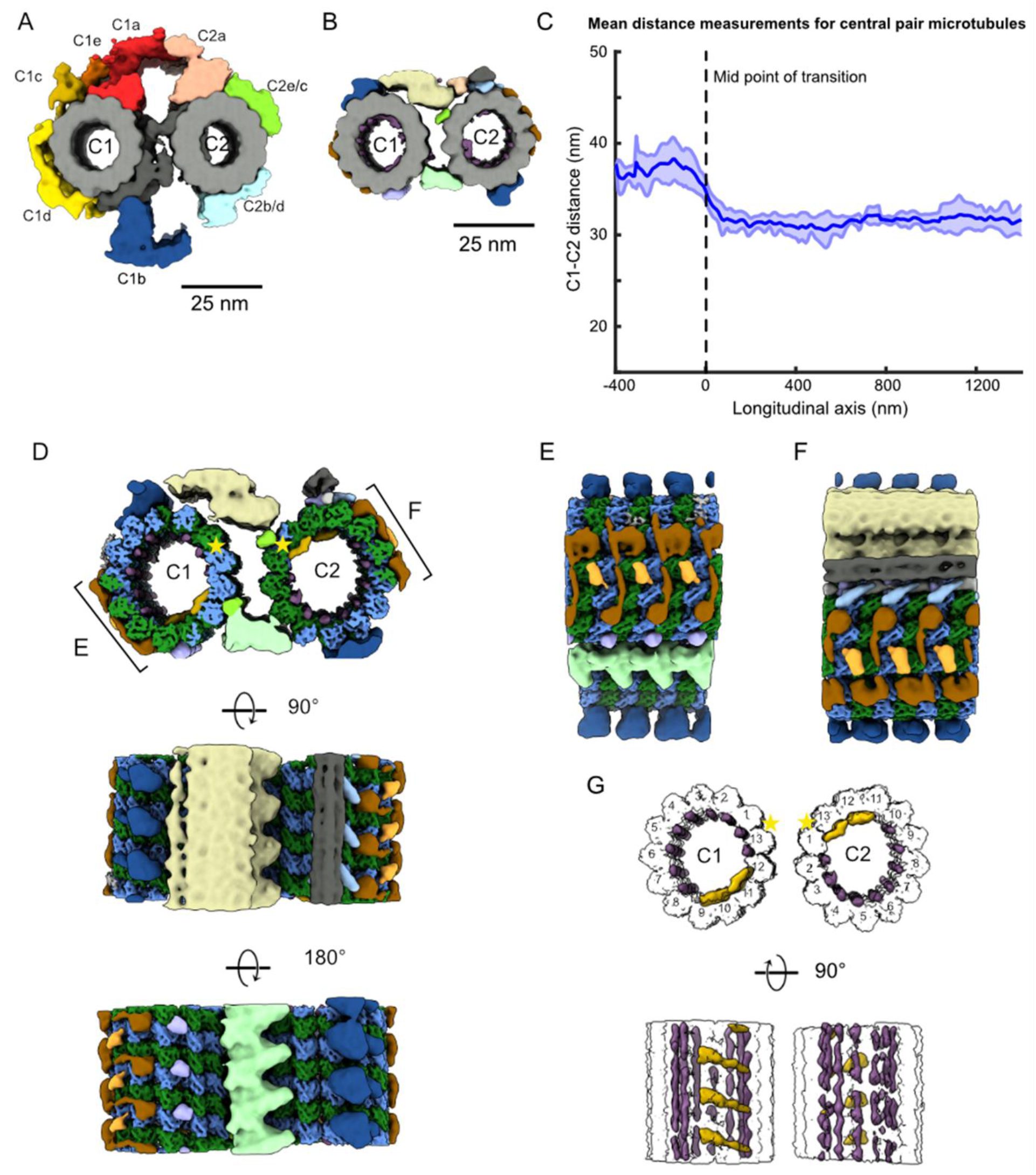
The tip CP has a different structure from the main CP. (A) Subtomogram average of the main CP. (B) Subtomogram average of the tip CP. In (A) and (B) structures lowpass filtered to 20 Å to be comparable. Microtubules are in grey, and projections have been coloured. (C) Graph of the distance between C1 and C2 microtubules. The reference on the x-axis is the middle of the narrowing zone. Average is shown in thick blue line. Standard deviation is shown in light blue. (D) Subtomogram average of the tip CP. Microtubules have been coloured according to tubulin subunits. Α-tubulin is coloured in green, and β-tubulin is coloured in blue. C1 is on the left and C2 is on the right. The yellow stars indicate the seams. (E) Side view of the tip CP as shown in (D). (F) Side view of the tip CP as shown in (D). (G) Tip CP rendering showing the MIPs in purple and yellow. The microtubules are in white and protofilaments are numbered from the seam. The yellow stars indicate the seams.

We then compared the structure of the main CP to the subtomogram average of the tip CP filtered to a similar resolution (Fig. 4B). Strikingly, the characteristic projections on the main CP are absent. The tip CP has smaller MAPs bound to the outside of C1 and C2. The spike densities observed in our tomograms (Fig. 1B) are shown in blue and localised to the top of C1- and the bottom of C2-microtubule (Fig. 4B, blue). The bridge between C1 and C2 is also different. The tip CP microtubules are linked by two densities: one at the top and one at the bottom (yellow and green densities, Fig. 4B).

When we visualised the subtomogram averages of the CP in the original tomograms (Fig. 1A), we observed clearly that the tip CP twists in an anti-clockwise direction. We measured the twist of the CP in the main part and compared it to that of the tip. We found that the tip CP twists almost three times more in the tip region: 0.72° every 8 nm versus 0.25° for the main CP.

The distance between C1 and C2 microtubules is significantly reduced in the tip CP. On average, C1 and C2 are 38.2 nm away from each other in the main region and 31.2 nm apart in the tip (Fig. 4C, S4D). The distance between C1 and C2 in the tip is similar to the distance between A-tubules (Fig. 3F). Interestingly, the narrowing zone seems consistent in all the tomograms. The distance between C1 and C2 always narrows over the length of around 100 nm. The shorter distance between C1 and C2 suggests that C1 and C2 are more tightly linked to each other in the tip region.

To increase the resolution of the tip CP and because the spike proteins observed in our tomograms repeated every 8 nm, we averaged particles picked every 8 nm. The resolution we obtained was about 9 Å which allowed us to distinguish between α- and β-tubulin subunits (Fig. 4D, S4E-F, Table S2). This allowed us to identify the seams on C1 and C2 microtubules (yellow stars). Both seams are facing each other similar to the main CP in *Chlamydomonas* (Gui, Wang et al. 2022, Han, Rao et al. 2022). C2 has more MAPs than C1 and is topped by a filament bundle which is perhaps similar to FAP7 found on top of C1 in the *Chlamydomonas* CP (Gui, Wang et al. 2022, Han, Rao et al. 2022).

A closer inspection of the tip CP structure shows that some MAPs bind laterally C1 and C2, contacting four protofilaments. These MAPs have a very similar shape suggesting they might be the same proteins (Fig. 4E and F, brown). We identified filamentous MIPs inside both C1 and C2 between adjacent protofilaments (Fig. 4G, purple). We also identified arc-MIPs making lateral contacts with 4 protofilaments in both C1 and C2 (Fig. 4G, yellow). Even though we cannot be certain about the repeating unit of the filamentous MIPs due to our imposed 8-nm periodicity averaging, our structure shows that all protofilaments are stabilised by either a longitudinal or arc-MIPs.

Surprisingly, we noticed that C1 and C2 are symmetric apart from the linker between the two microtubules unlike in the clear difference in main-CP region. The MIPs and MAPs on C1 overlay with the MIPs and MAPs on C2 when it is rotated 180° around the longitudinal axis (Fig. 4D to G) suggesting that the same proteins assemble in and around both C1 and C2.

### *FAP256A/B-KO* mutants lack the ciliary cap

The current resolution of our subtomogram averages does not allow us to identify proteins *de novo*. Only a few proteins are known to localise to the ciliary tip in *Tetrahymena* including FAP256 (A and B), CHE-12 (A and B) and ARMC9 (A and B). We decided to use cryo-ET to study the cilia of cells lacking FAP256 (*FAP256A/B-KO*) because of the lack of ciliary cap complex reported (Louka, Vasudevan et al. 2018). Analysis of the tomograms obtained revealed various defects in the tips. First, we observed that the CP cap was absent in all our tomograms while cilia had a normal overall shape (Fig. 5A, S5A). The filaments going inside the CP microtubules were still present (Fig. 5B, S5A, Fig. 2B). Although their molecular identities remain unknown, the similarity to the plug observed at the end of the A-tubules suggests that they are composed of the same proteins. Looking closely at the tip in *FAP256A/B-KO* mutants, we saw that the CP microtubules end with slightly curved protofilaments that contact the membrane directly (Fig. 5B and C). The curvature of the protofilaments seems higher than that of sperm singlets when they reach the membrane (Zabeo, Heumann et al. 2018, Zabeo, Croft et al. 2019). We did not measure the curvature of individual protofilaments as we cannot resolve them in the WT tomograms due to the ciliary cap and therefore cannot compare the curvature to a control. Although microtubules appeared flared at their ends, we did not observe any tapered ends.

**Figure 5.**
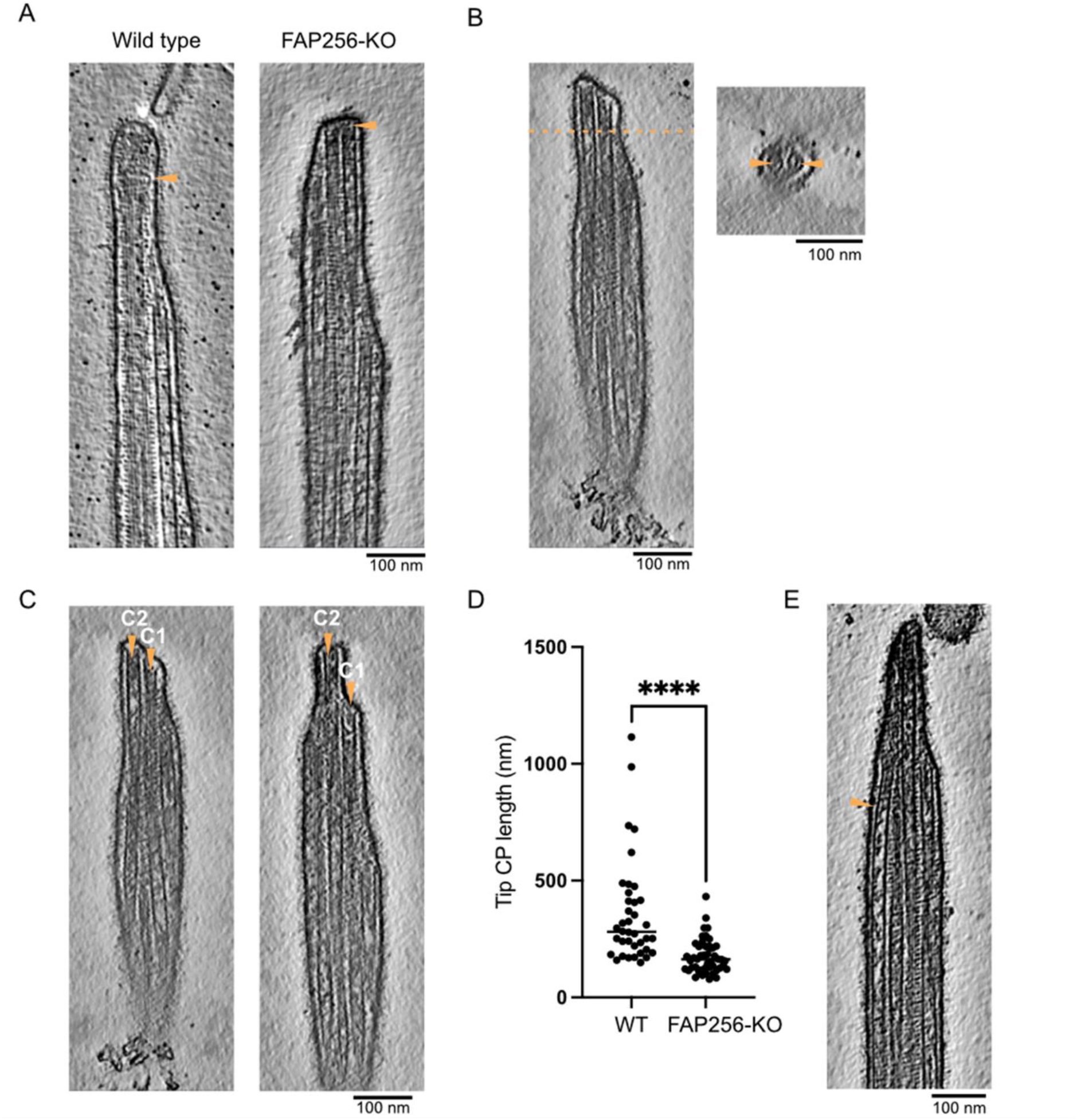
FAP256-KO ciliary tips show multiple structural defects. (A) Slices of *WT* and *FAP256A/B-KO* tomograms. The arrow points to the end of the CP microtubules to highlight the lack of cap in mutant tomograms. (B) Tomogram slice of *FAP256A/B-KO* cilium along with cross-section (dotted line). The arrows indicate the densities inside the CP microtubules that correspond to the plugs. (C) Tomogram slices of *FAP256A/B-KO* cilia. The arrows indicate the end of the CP microtubules. (D) Quantification of tip CP length in *WT* and *FAP256A/B-KO* cilia. Each dot represents one CP. **** p-value<0.0001, Mann-Whitney test. (E) Tomogram slice of a *FAP256A/B-KO* mutant cilium. The arrow points to a bent singlet A-tubule.

While analysing our tomograms, we also observed that often, C1 and C2 had different lengths (Fig. 5C) while they would typically have the same length in WT cilia. In the two examples shown, C2 is longer than C1. These defects in CP microtubules prompted us to measure the length of the tip CP, that is the CP portion covered by the spike protein (Fig. 5D). Indeed, the tip CP was significantly shorter in *FAP256A/B-KO* than in *WT*. These results agree with a previous study (Louka, Vasudevan et al. 2018).

To verify whether the CP in *FAP256A/B-KO* had defects, we used subtomogram averaging to obtain a structure of the tip CP. At the same resolution, both *WT* and *FAP256A/B-KO* tip-CP were identical (Fig. S5B, Table S2). The A-tubules appeared identical to WT. The plugs at their ends were also present (Fig. S5C). Interestingly, we observed that multiple A-tubule singlets were curved towards the CP when they are normally straight in WT cells (Fig. 5E, Fig. S5D, Fig. 3F).

As the ciliary cap complex was absent from *FAP256A/B-KO* cilia, we next sought to identify the proteins found in this complex. To this end, we carried out mass spectrometry to identify the missing proteins in mutant cilia (Fig. 6A). Both FAP256A and FAP256B was indeed absent, confirming that our assay worked. We identified six other proteins that were absent from *FAP256A/B-KO* cilia (Table 1). Interestingly, FAP256 was identified in a screen for tip-specific proteins in *Chlamydomonas* (Satish Tammana, Tammana et al. 2013) among other proteins. However, none of the proteins that we identified as absent from *FAP256A/B-KO* were found in the previously published screen. This observation is likely due to the fact that we did not find homologs for the proteins we identified. The proteins that are absent from *FAP256A/B-KO* include CCDC81 which is found in humans but not *Chlamydomonas*. It is found at centrosomes in human cells although not found in human primary cilia (Firat-Karalar, Sante et al. 2014). The predicted structure of CCDC81 protein (TTHERM_00500930) contains an N-terminal CCDC81 domain and several EF-hand domain pairs in the C-terminus (Fig S6A). The N-terminal CCDC81 domain aligns well with that of IJ34, another CCDC81 homolog, recently identified as a B-tubule MIP binding on the surface of the A-tubule at the inner junction (Kubo, Black et al. 2022). However, when overlaid with α-tubulin, parts of the microtubule-binding domain of IJ34 are missing in CCDC81. Despite this, we believe it still likely binds microtubules (Fig. S6B). The EF-hand domains of the tip-CCDC81 suggests it might be involved in calcium regulation.

**Table 1.** List of proteins identified as absent from *FAP256A/B-KO* cilia compared to WT shown in Fig. 5A.

| Uniprot ID | Name | Amino acids | Human homolog? | Known localisation? |
| --- | --- | --- | --- | --- |
| I7MLF6 | EF hand protein | 1766 | CCDC81 | centrosome |
| Q22US8 | DUF4476 domain-containing protein | 798 | no |  |
| I7MKU9 | Transmembrane protein | 1730 |  |  |
| Q24IJ6 | Tetratricopeptide repeat protein | 519 | TPR repeat protein 28 | microtubule organising centre |
| Q23FN3 | FAP256 | 872 | CEP104 | cilia and centriole |
| I7LVP2 | Divergent AAA domain protein | 273 | Li 173 or SLFNL1 | sperm cells |
| I7MH51 | CLASP_N domain-containing protein | 474 |  |  |

**Figure 6.**
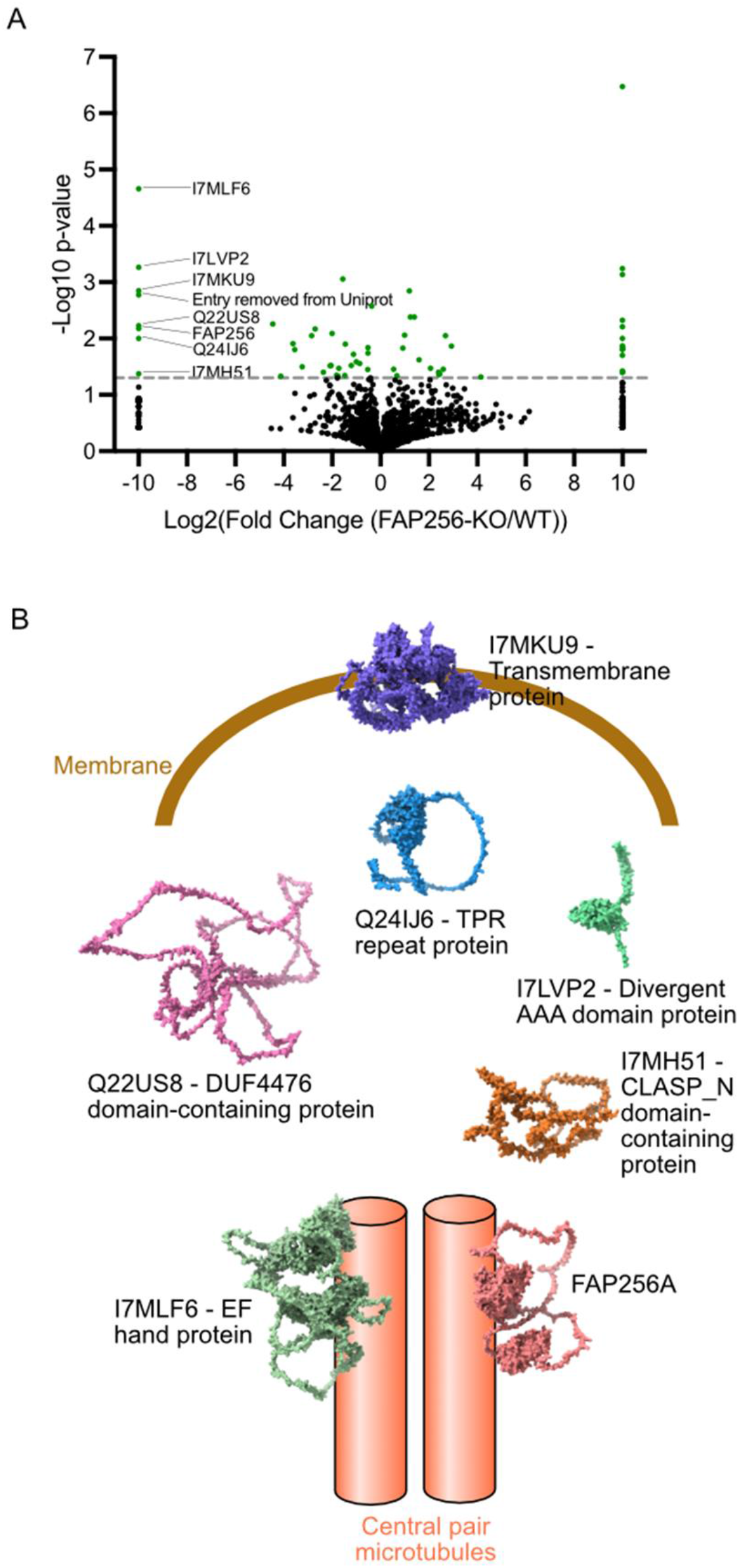
The ciliary tip complex proteins are long and flexible. (A) Volcano plot of the peptides identified in *FAP256A/B-KO* vs *WT* cilia. The proteins absent in *FAP256A/B-KO* are listed on the left-hand side of the graph. (B) Surface representation of the proteins identified in (A) and predicted with AlphaFold 2. Proteins predicted to bind microtubules are represented close to the CP microtubules.

Ca^2+^ ions have been shown to change the direction of swimming in *Tetrahymena* (Izumi and Miki-Noumura 1985, Marumo, Yamagishi et al. 2021). Proteins with EF-hand pair domain were also recently identified as doublet MIPs in *Tetrahymena* such as CFAP115, Rib57 and Rib22 (Kubo, Black et al. 2022). It is therefore possible that the N-terminal of CCDC81 binds the outside of the microtubule while the C-terminal EF-hand domains bind the inside of the microtubules. We also identified SNFL1 which is likely expressed in mammalian sperm cells. In addition, sperm-tail PG-rich repeat protein and tubulin-tyrosine ligase were absent although not significant. Recently, sperm-tail PG-rich proteins were identified as MAPs binding both insides and outside microtubules (I7M2G0 and Q24GM1) (Kubo, Black et al. 2022). The other proteins that we identified were predicted to be long and disordered and have not been characterised previously. Based on their predicted domains, we identified one protein as membrane-binding (I7MKU9, Fig. 6B) and two proteins as microtubule-binding proteins (FAP256 and CCDC81, Fig. 6B). The other proteins are likely to act as linker proteins that link the membrane to the CP microtubules. All the missing proteins in *FAP256A/B-KO* mutant are present in the proteome of demembranated WT doublet microtubules, in which the ciliary cap is still present (Kubo, Yang et al. 2021). We also listed the proteins that had a fourfold reduction in *FAP256A/B-KO* cilia (Table S3). These proteins include a predicted kinesin (I7LVY1) that is likely to bind the CP microtubules in the cap complex. Based on these results, we believe these proteins are therefore likely to be part of the ciliary cap complex.

## Discussion

### The architecture of the ciliary tip differs from the rest of the axoneme

Cryo-ET allowed us to study in detail the overall architecture of the motile ciliary tips. Measuring the starting and ending points of the A-tubules and their lengths revealed a great variety. This differs from the rest of the axoneme, which has molecular rulers that regulate the spacing between the various subcomplexes making up the axoneme. Recently, three proteins were identified as regulators of the length of A- and B-tubules. FAP256 promotes the elongation of A-tubules while CHE-12 and ARMC9 promote the elongation and depolymerisation of B-tubules (Louka, Vasudevan et al. 2018). The outside of the doublet microtubule has a periodicity of 96 nm that is regulated by the coiled coil proteins CCDC39 and CCDC40 (Oda, Yanagisawa et al. 2014). It is likely that the availability of these proteins regulates the assembly of the last radial spokes and dynein arms. We have shown that B-tubules are often not complete from the inner junction to the outer junction suggesting that the inner junction is less stable or incompletely assembled. Isolated doublets from *Tetrahymena* were seen to be less stable at the inner junction (Khalifa, Ichikawa et al. 2020). The inner junction is made up of multiple proteins with CFAP52 as an interaction hub. It is likely that this protein along with ARMC9 regulate the disassembly of the B-tubule. B-tubules are highly glutamylated (Lechtreck and Geimer 2000). Glutamylation and other microtubule post-translational modifications might also play a role in the termination of the B-tubules.

Interestingly, both A-tubules and CP microtubules contain non-periodic MIPs. MIPs of similar size have been identified in neurons and primary cilia (Garvalov, Zuber et al. 2006, Bouchet-Marquis, Zuber et al. 2007, Kiesel, Alvarez Viar et al. 2020, Foster, Ventura Santos et al. 2022). The identities of these MIPs remain a mystery but some could be microtubule-modifying enzymes such as acetyltransferases. They might also be important for the stability of the microtubules, but we think this is an unlikely scenario for two reasons: one, because the CP microtubules are already stabilised by filamentous and arc-MIPs and two, because the other microtubule-stabilising proteins found in cilia have a constant periodicity. Intriguingly, our subtomogram average of the A-tubule does not show any stabilising protein, raising the question of how these are stabilised. In human, bull, pig and horse sperm cells, SPACA9 provides stability to the A-tubule by forming a helix inside the lumen (Zabeo, Heumann et al. 2018, Leung, Roelofs et al. 2021, Gui, Croft et al. 2022, Leung, Roelofs et al. 2022). This difference likely reflects the role of the ciliary tip. The tips of sperm cells might need to be more resistant to forces as opposed to the tips of *Tetrahymena*.

Microtubules are dynamic and unstable at their plus ends. They must therefore be stabilised in cilia. The plugs seen at the ciliary tip are likely to play this role. Proteins stabilising the plus ends of centriolar microtubules have been studied previously. CEP110 and CEP97 act as a plug that can be visualised *in vitro* using tomography (Ogunmolu, Moradi et al. 2022).

IFT trains convert from anterograde to retrograde at the tips. The observation that anterograde trains walk on the B-tubules raises the question of how these trains reach the ciliary tip when the B-tubules end. The IFT train-like densities we observe seem more similar to that of anterograde than retrograde trains. Anterograde trains have already been observed on the A-tubules in primary cilia (Kiesel, Alvarez Viar et al. 2020). In our tomograms, we did not observe retrograde-train-looking particles (Jordan, Diener et al. 2018). Whether the densities we observed are active anterograde trains on A-tubules remains to be determined.

### The CP changes morphology in the tip

When reaching the ciliary tip, the CP is entirely reshaped. At our resolution, we were not able to identify densities shared between the main and tip portions of the CP. The mechanism that regulates the lengths of the two CP segments remains to be discovered. The different structures suggest a dual role for the CP. In the main portion of the axoneme, the CP acts as a signal transducer between the radial spokes and its large projections (Oda, Yanagisawa et al. 2014). In the tip, the CP may act as a strengthening rod for the ciliary tip.

The striking symmetry observed between C1 and C2 suggests that the same proteins assemble around the two CP microtubules. One exception is a small protein located at the seam of C2. This protein must recognise the non-canonical interaction between shifted tubulin dimers. The density for this protein overlays very well with the one found on C1 between protofilaments 11 and 12 (Fig. 3D). The protein on C1 is likely to be different as the C1 seam is located elsewhere. The seams facing each other is consistent with the seam-centric model for the assembly of the CP (Gui, Wang et al. 2022). Although the seams for the main CP could not be identified due to insufficient resolution, we believe they are located at similar positions to the tip CP, which is also consistent with the *Chlamydomonas* CP (Gui, Wang et al. 2022, Han, Rao et al. 2022).

This study raises questions on the mode of assembly of the CP. There is evidence suggesting that the CP assembles in a unique manner, distinct from the peripheral axonemal microtubules. In *Chlamydomonas* dikaryons that repair their CP under conditions of genetic complementation, the CP initially assembles as short segments that are positioned away from either the basal body or the ciliary distal end. Possibly, these fragments later fuse to form the CP (Lechtreck, Gould et al. 2013). Whether the CP assembles as multiple fragments during the normal course of ciliary elongation is still unknown. In *Tetrahymena*, a recent study showed that the CP was present in the tips of short growing cilia however its precise structure and elongation mechanism remain undetermined (Reynolds, Phetruen et al. 2018). Regardless of the mode of CP assembly, it remains to be determined how the boundary between the main and tip segment of the CP is positioned during ciliary assembly.

### The tip is a highly stable entity

This distal region must be very stable to resist the various forces at the tips of *Tetrahymena* cilia. Various lines of evidence point to that direction. First, we see that the CP microtubules are closer to each other, indicating that the bridge has shortened to restrict movement between the two microtubules. Second, the spike proteins binding to the outside of the CP microtubules repeat every 8 nm as opposed to 16 or 32 nm for the main CP. Third, the conformation of MAPs binding on the outside of C1 and C2 is such that multiple protofilaments are engaged and therefore strengthens the lateral interaction of tubulin. Microtubule interactions are stronger in the longitudinal direction (VanBuren, Odde et al. 2002). Lateral MIPs and MAPs are likely to be important to stabilise microtubules while cilia are moving in different directions. These different MAPs with a shorter periodicity might explain the higher twist seen in the ciliary tip. Fourth, we observe links between A-tubules and between A-tubules and the membrane in the tip that contribute to their stability.

### FAP256 contributes to the stability of the ciliary tip

The main two defects we observed in *FAP256A/B-KO* mutants are bending of singlets and sliding of one of the CP microtubules compared to the other. The CP microtubules are capped by the tip complex in WT cells which is absent in mutant cells. FAP256 therefore might prevent the sliding of CP microtubules along with the proteins of the tip complex which likely serve as an anchor for the CP microtubules to the membrane. The difference in length observed might be due to a microtubule that is longer than the other. We identified CCDC81 and several others as CP cap proteins. As it contains domains that are predicted to bind inside and outside the microtubule, it is possible that is acts as a regulator of microtubule polymerisation. From our tomograms, it is unclear whether the links between A-tubules or between A-tubules and membrane are absent, but the bending of the A-tubules observed in our *FAP256A/B-KO* tomograms suggests that some link is missing. We therefore propose that FAP256, along with its interacting-proteins contributes to stabilising the ciliary tip microtubules by preventing sliding and bending of these microtubules (Figure 7). C2 in *Chlamydomonas* can slide with regards to C1 in the medial region of the cilium. The sliding distance measured was up to 24 nm (Han et al., 2022). Our model explains the observed reduced functionality of motile cilia caused by the loss of FAP256 in *Tetrahymena* (Louka, Vasudevan et al. 2018). It is currently unclear whether the same sliding occurs in other species and in the tip of *Tetrahymena thermophila*. The ciliary cap is possibly essential to maintain the association of C1 and C2 during sliding.

**Figure 7.**
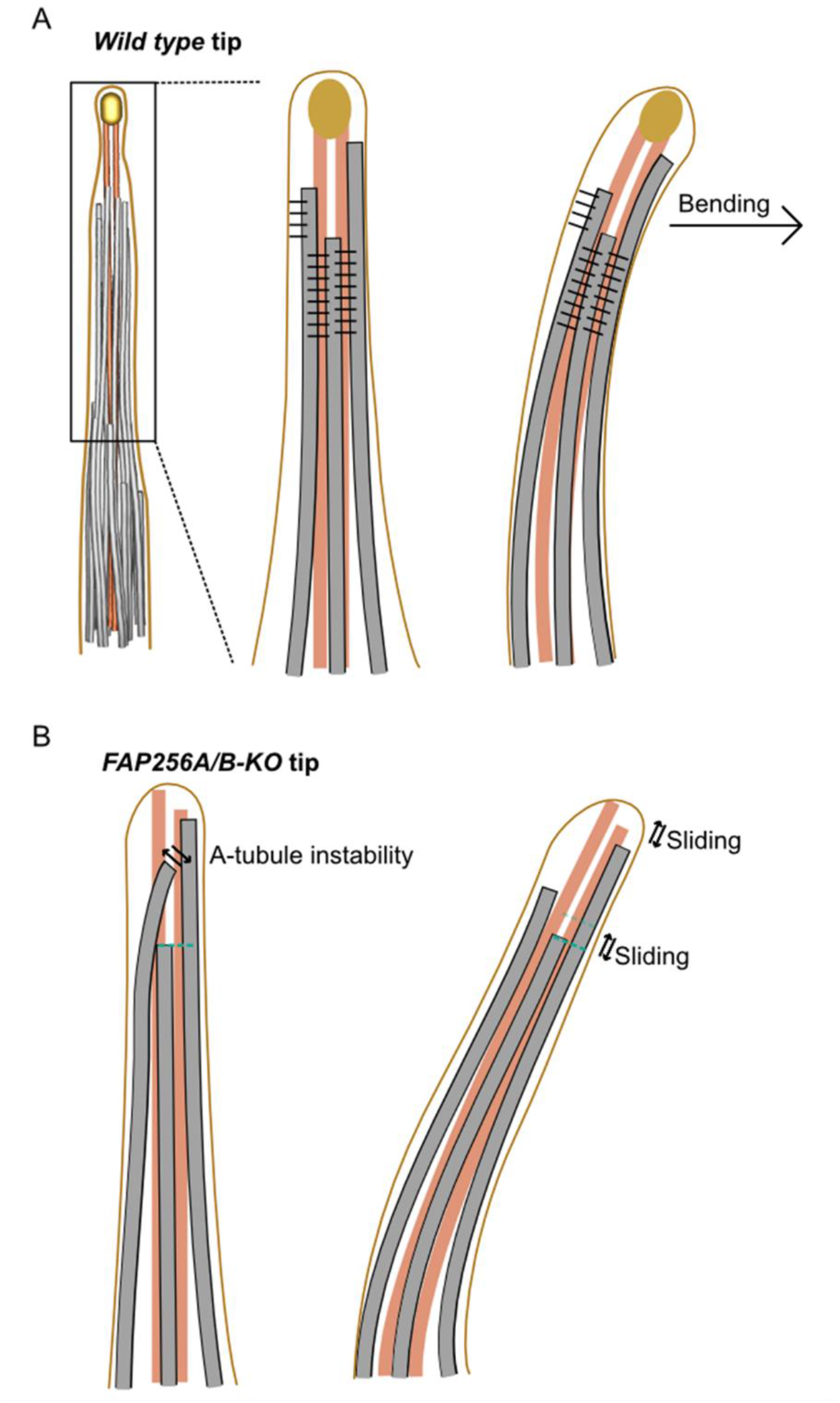
The A-tubule links stabilise the A-tubules and restrict sliding. (A) Diagram of WT cilium with the A-tubule and membrane links drawn in black in straight and bending cilium. (B) Diagram of *FAP256A/B-KO* mutant cilia without the A-tubule links showing bent A-tubules and sliding A-tubules and CP microtubules during bending.

This study represents the most detailed description of the ciliary tip of *Tetrahymena thermophila* to date. Future studies will be needed to identify the proteins that bind to the CP microtubules and shed light on their precise functions.

## Materials and methods

### Cell culture

All *Tetrahymena* strains (*CU428* wild-type, *FAP256A/B-KO* mutant) used in this study were grown in SPP media (Williams, Wolfe et al. 1980) in a shaker incubator at 30°C and 150 rpm. Cultures were harvested at an OD_600_ of approximately 0.6. Refer to (Black, Dai et al. 2021) for further details.

### Cilia purification

*Tetrahymena* cells were harvested and deciliated as previously described (Black et al., 2021). Briefly, a 2 L cell culture was harvested by centrifugation at 2,000 g for 10 minutes at 23°C. Whole cells were resuspended in fresh SPP and adjusted to a total volume of 24 mL. Dibucaine (25 mg in 1 mL SPP) was added, and the culture was gently swirled for 45 seconds. To stop the dibucaine treatment, 75 mL of ice-cold SPP supplemented with 1 mM EGTA was added and the dibucaine-treated cultures were centrifuged for 15 minutes at 2,500 g and 4°C. The supernatant (cilia) was collected and centrifuged at 25,000 g for 45 minutes at 4°C. The pellet (cilia) was gently washed and resuspended in Cilia Wash Buffer (50 mM HEPES at pH 7.4, 3 mM MgSO_4_, 0.1 mM EGTA, 1 mM DTT, 250 mM sucrose) before snap freezing in liquid nitrogen for long-term storage. Cilia were resuspended in Cilia Final Buffer (50 mM HEPES at pH 7.4, 3 mM MgSO_4_, 0.1 mM EGTA, 1 mM DTT, 0.5% trehalose) before grid freezing. NP-40 Alternative (Millipore Sigma #492016) was added to a final concentration of 1.5% for 45 minutes on ice to remove the membrane. Axoneme were then centrifuged and resuspended in Cilia Final Buffer.

### Cryo-ET preparation

The axonemes for cryo-ET were cross-linked with glutaraldehyde (final concentration 0.15%) for 30 min on ice and quenched with 35 mM Tris pH 7.5. Axoneme concentration was approximately 3.6 mg/mL when mixed with 5 (Cytodiagnostics) or 10 (Aurion) nm gold beads in a 1:1 ratio for a final axoneme concentration of 1.8 mg/mL. Crosslinked axoneme sample (4 μl) was applied to negatively glow-discharged (10 mA, 10 s) C-Flat Holey thick carbon grids (Electron Microscopy Services #CFT312-100) inside the Vitrobot Mk IV (Thermo Fisher) chamber. The sample was incubated on the grid for 45 seconds at 23°C and 100% humidity before being blotted with force 0 for 8 seconds and plunge frozen in liquid ethane.

### Tilt series acquisition

Tilt series were collected at 42,000x and 26,000x magnifications using the grouped dose-symmetric scheme from -60 to 60 degrees with an increment of 3 degrees (136, 64 and 33 tilt series of demembranated *WT*, membranated *WT* and *FAP256A/B-KO*, Table S1). The defocus for each tilt series ranged from -2 to -6 μm. The total dose for each tilt series was 164 e_-_ per Å_2_. For each view, a movie of 10-13 frames was collected. The pixel size at 42,000x and 26,000x were 2.12 Å and 3.37 Å, respectively. Frame alignment was done with Alignframes (Mastronarde and Held 2017).

Tomograms were reconstructed using IMOD (Kremer, Mastronarde et al. 1996). Aligned tilt series were manually inspected for quality and sufficient fiducial presence. Batch reconstruction was performed in IMOD.

### Subtomogram Averaging

CTF estimation of the tomogram was done with WARP (Tegunov and Cramer 2019). The picking of the CP and A-tubules was done using IMOD by tracing line along the microtubules (Kremer, Mastronarde et al. 1996). The subtomogram particles were then picked at 8-nm (tip-CP and A-tubules) or 16-nm (base-CP) intervals along the traced lines. The pre-alignment of the 4-times binned 8-nm repeating unit of the tip-CP of *WT* and *FAP256A/B-KO* mutants (11,534 and 4,407 subtomograms respectively) was done using a set of Dynamo (Castaño-Díez, Kudryashev et al. 2012) scripts for filament alignment (https://github.com/builab/dynamoDMT). Then, the alignment parameters were converted into Relion 4.0 format. Refinement of the subtomogram averages was done with the Relion 4.0 pipeline (Kimanius, Dong et al. 2021).

After importing the prealigned subtomogram into Relion 4.0, the following workflow was followed: (i) Generation of pseudosubtomogram; (ii) Refinement of the subtomograms; (iii) CTF Refinement; (iv) Frame Alignment; (v) Pseudo-tomogram generation and (vi) Refinement of the polished subtomograms (Fig. S4A).

The resolutions for the 8-nm repeating unit of the tip-CP of *WT* and *FAP256-KO* DMT are 9 and 23 Å, respectively.

For the base-CP, after obtaining the 16-nm averages, we performed a 3D classification into 32-nm repeat of the base-CP. The resolution of the base-CP is 23 Å.

### Visualisation/Segmentation

For denoising, tomograms were CTF deconvolved and missing wedge corrected using IsoNet (Liu, Zhang et al. 2022). The IMOD model of the ciliary tip was generated from both automatically converting the aligned subtomogram coordinate (microtubules, tip complex) and manually tracing the tomograms (MIPs, plugs and spacers). Membrane segmentation was done on the denoised tomograms using TomoSegMemTV (Martinez-Sanchez, Garcia et al. 2014). For visualisation of the subtomogram particles inside the tomogram coordinate (Fig. 1A), Subtomo2Chimera (10.5281/zenodo.6820119) was used.

For subtomogram averages, surface rendering, segmentation and fitting were done using ChimeraX (Pettersen, Goddard et al. 2021).

### Measurement of CP twist

To measure the CP twist in the base and the tip, the same subtomogram average of the CP was loaded twice into ChimeraX. One average was then shifted 8-nm or 16-nm from the other average for the tip and the base CP respectively. The ChimeraX *fitmap* function was then used to fit the averages together and the twist angle was read out from the alignment.

### Measurement of distance

The distance between C1 and C2 microtubules was done through a custom Matlab script. These microtubules were first picked independently in the tomograms containing the narrowing zone. Subtomogram averaging of each microtubule was then performed using particles picked every 16 nm to properly centre the C1 and C2 microtubules. Using the centre of each particle, a spline line was extrapolated along the C1 microtubule and the shortest distance from each point of C2 microtubule to C1 microtubule line was measured (Fig. S4D). The distance measured between C1 and C2 from different tomograms was plotted on a graph (Fig.4C). The inflection point (midpoint) was identified automatically by determining the points with the most negative slope. Each CP graph was shifted such that the midpoint was situated at length 0. A mean graph was finally generated from multiple tomograms.

A similar analysis was carried out to measure the distance between A-tubule pairs.

### Mass spectrometry

Mass spectrometry was performed as described in a previous publication (Khalifa, Ichikawa et al. 2020). Approximately 25-30 μg of membranated cilia was loaded on the SDS-PAGE gel. The electrophoresis run was terminated before the proteins entered the resolving gel. The band containing all proteins was cut out and subjected to in-gel digestion. After that, the peptides were separated on a Dionex Ultimate 3000 UHPLC and loaded onto a Thermo Acclaim Pepmap (Thermo, 75 μm ID × 2 cm with 3 μm C18 beads) precolumn and then onto an Acclaim Pepmap Easyspray (Thermo, 75 μm × 25 cm with 2 μm C18 beads) analytical column. The peptides were separated with a flow rate of 200 nl/min with a gradient of 2-35% solvent (acetonitrile containing 0.1% formic acid) over 2 hours. Peptides of charge 2+ or higher were recorded using a Thermo Orbitrap Fusion mass spectrometer. The data was searched against *Tetrahymena thermophila* protein dataset from UniProt (https://www.uniprot.org/). MS data were analysed by Scaffold_4.8.4 (Proteome Software Inc.). Statistically significant proteins missing in *FAP256A/B-KO* mutant compared to *wild type* were identified as cap complex candidates.

### AlphaFold prediction

AlphaFold prediction of the cap complex protein candidates were downloaded from AlphaFold Protein Structure Database (https://alphafold.ebi.ac.uk/) except for I7MLF6 which was predicted using the COSMIC_2_ web platform (Cianfrocco, Wong-Barnum et al. 2017).

### Data Availability

The data produced in this study are available in the following databases:

- Cryo-EM map of the base-CP, tip-CP, doublet microtubule and A-tubule: EMDB EMD-XXXX, EMD-YYYY, EMD-ZZZZ, EMD-XYXY
- Cryo-EM map of the tip-CP from *FAP256A/B-KO* mutant: EMDB EMD-XXXX,

The mass spectrometry data of *FAP256A/B-KO* and *CU428* (WT) is available in DataDryad at XXXX.

The Matlab code for subtomogram averaging in Dynamo and distance analysis is available at https://github.com/builab/dynamoDMT

## Authors contributions

Conceptualisation: KHB. Methodology: TL. Formal analysis: TL, MT. Investigation: TL, MT, CB, MPV, KHB. Resources: JG, KHB. Writing–original draft: TL, KHB. Writing– review & editing: TL, KHB, CB, JG.

## Acknowledgments

We thank all the members of the Bui lab for critical reading of the manuscript. We thank Drs. Mike Strauss, Kaustuv Basu, Kelly Sears (Facility for Electron Microscopy Research at McGill University) for helping with data collection. We thank Amy Wong, Lorne Taylor and Jean-François Trempe (RI-MUHC Proteomics Platform) for their help with mass spectrometry. KHB is supported by the grants from Canadian Institutes of Health Research (PJT-156354) and Natural Sciences and Engineering Research Council of Canada (RGPIN-2022-04774). JG is supported by NIH grant R01GM135444.

## Supplementary figures

**Supplementary Figure 1.**
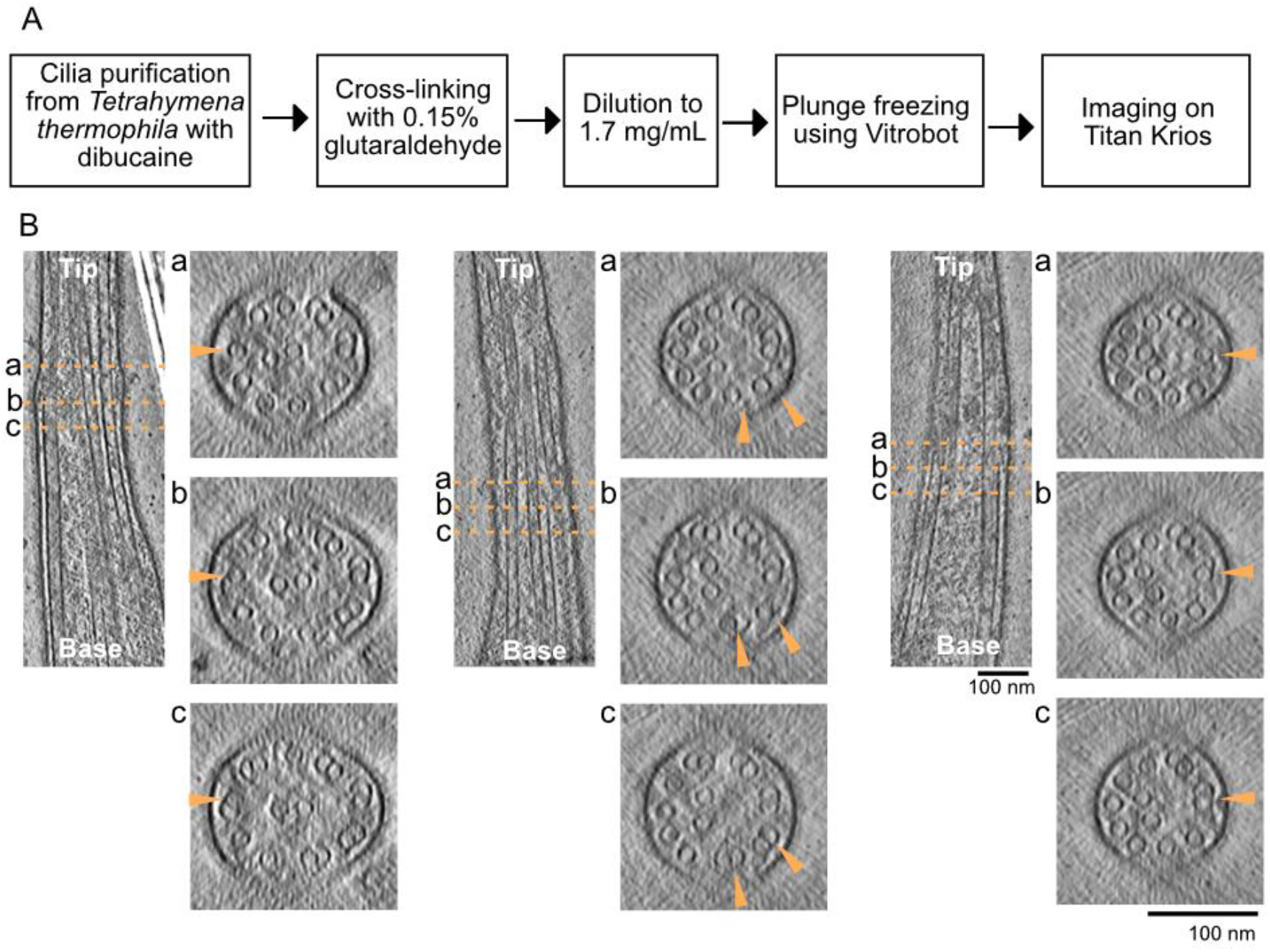
(A) Diagram of cilia preparation workflow. (B) Longitudinal slices through tomograms and corresponding cross-sections as in Fig. 1E. Arrows point to disassembling doublets. Scale bar 100 nm.

**Supplementary Figure 2.**
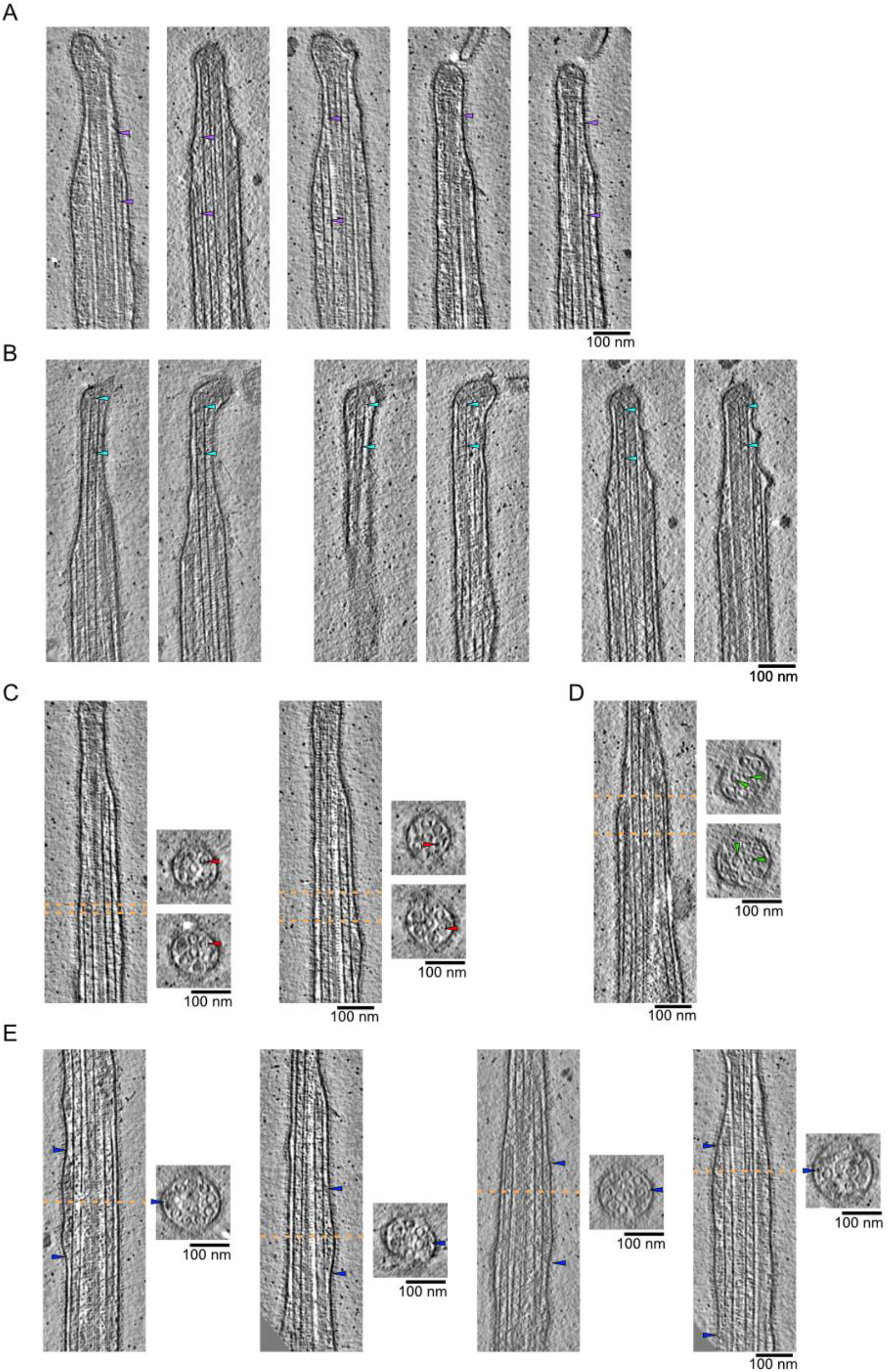
(A and B) Examples of filamentous plugs in A-tubules (A) and CP microtubules (B). Arrows point to the extremities of the filamentous plugs. (C, D, E) Examples of MIPs found in A-tubules (C), CO microtubules (D) and IFT train-like particles (E). Longitudinal slices along tomograms. Dotted lines indicate cross-sections. Scale bars: 100nm.

**Supplementary Figure 3.**
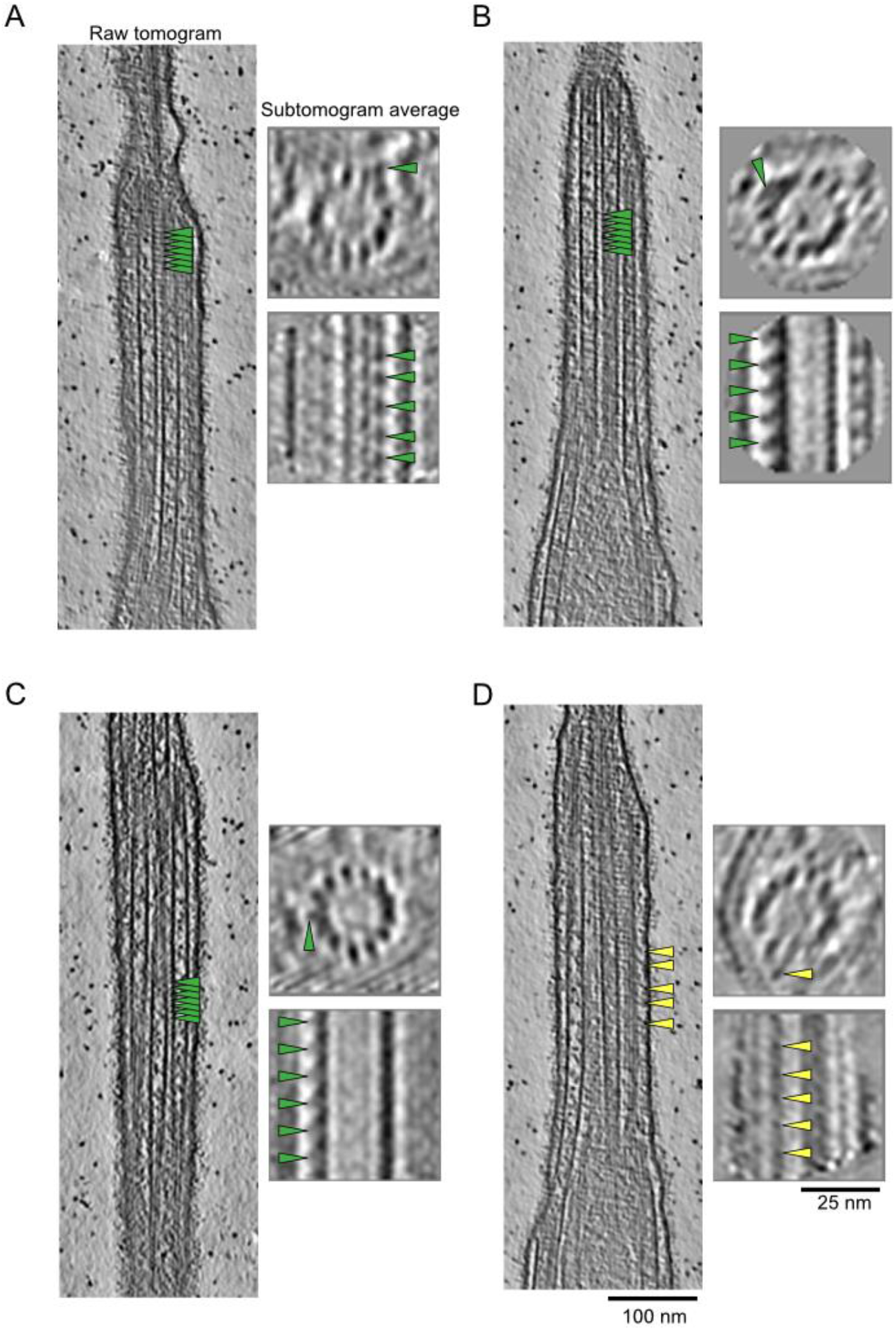
Examples of A-tubule links (A-C) and membrane links (D) as in Fig. 3 (B-E). Representative raw tomogram longitudinal slices along with cross and longitudinal sections of subtomogram averages of individual A-tubules. Green and yellow arrows point to A-tubule and membrane links respectively.

**Supplementary Figure 4.**
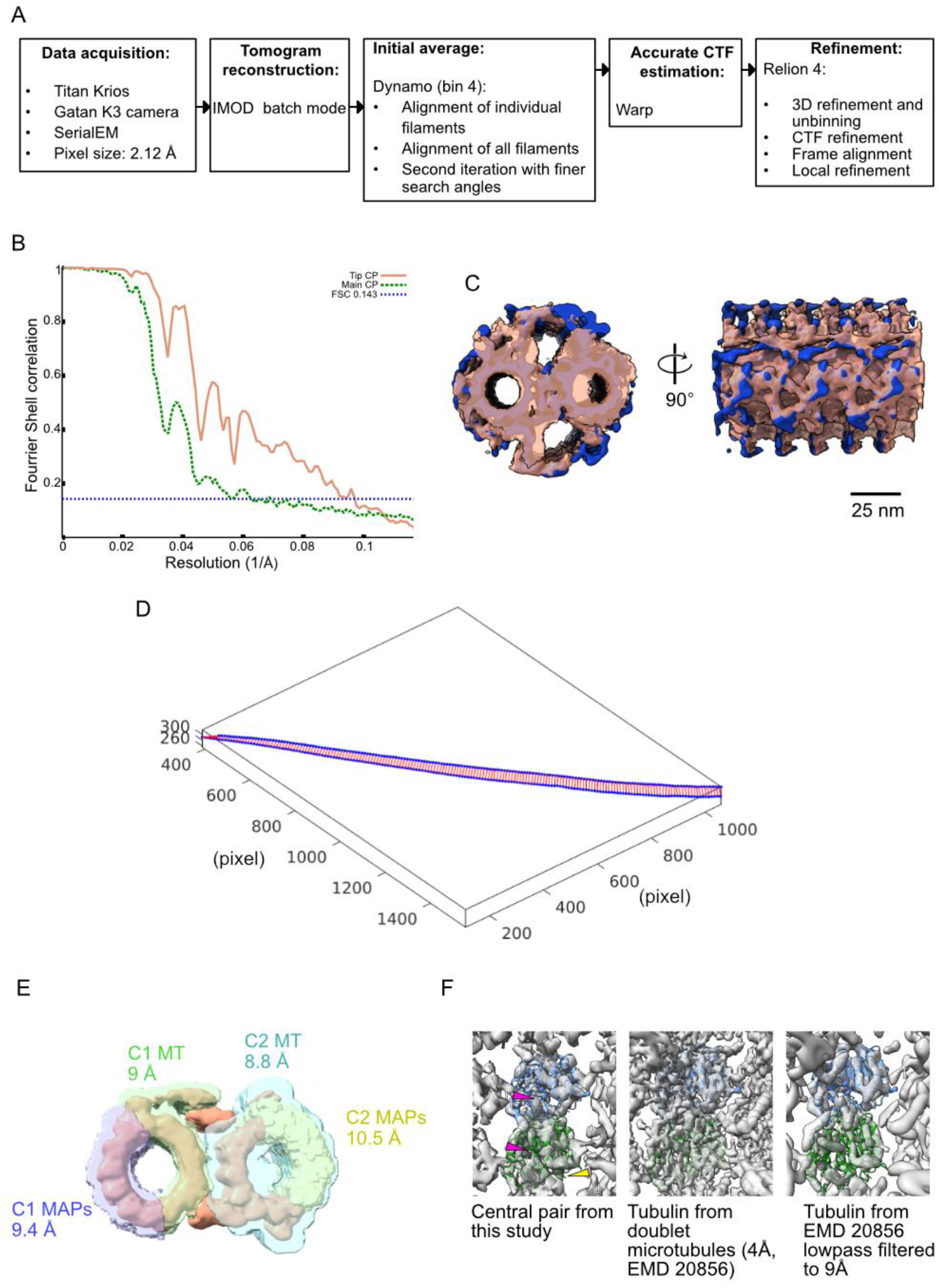
(A) Diagram of the subtomogram averaging workflow used in this study. (B) Fourier Shell Coefficient (FSC) curves of the Tip CP and main CP structures. (C) Front and side view of *Tetrahymena* main CP (this study, dark salmon, transparent) overlaid with sea urchin CP (EMD 9385, navy blue). (D) Example of measurement of the distance between both CP microtubules along the length of a tomogram. The blue dots represent the coordinates of the centre of a microtubule subtomogram. The red lines represent the shortest distance between two blue points. (E) Masks used to refine the structure of the tip CP in this study. The resolution obtained after masking is indicated on the figure for each mask. (F) Fitting of a tubulin dimer in the electron density map of the subtomogram average of the tip CP. Red arrows point to the S loop which is long for α-tubulin and short for β-tubulin. α-tubulin is coloured in green and β-tubulin is coloured in blue. Yellow arrow points to K40 loop. For comparison, the same tubulin dimer was fitted in a map at 4 Å resolution which was then lowpass filtered to 9 Å.

**Supplementary Figure 5.**
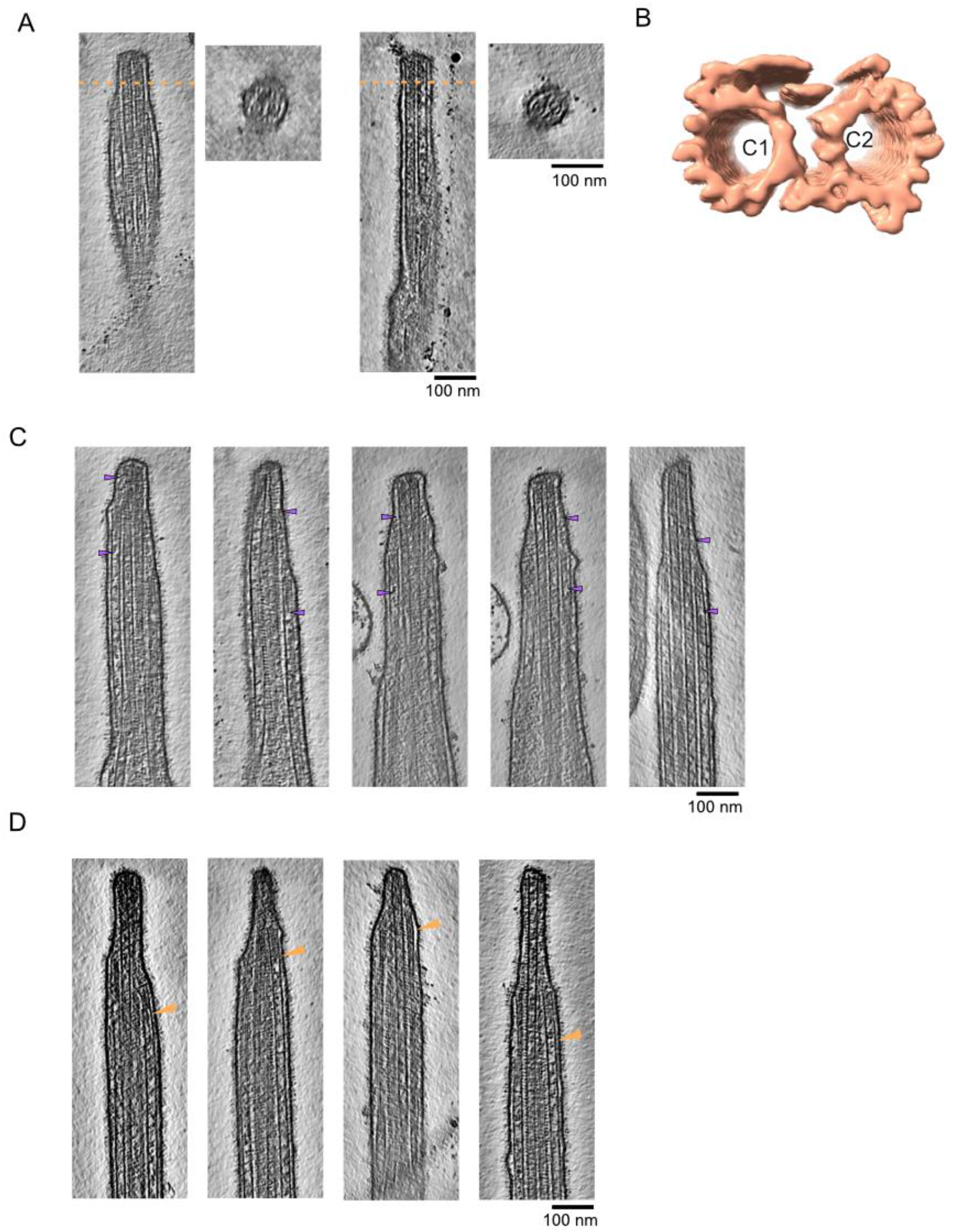
(A) Slices of *FAP256A/B-KO* tomograms along with cross-section (dotted line) showing examples of the filamentous plugs in both CP microtubules. (B) Subtomogram average of *FAP256A/B-KO* tip CP. (C) Gallery of A-tubule plugs. The arrows point to the extremities of the plugs. (D) Gallery of bent A-tubules indicated by arrows. Scale bars: 100nm.

**Supplementary Figure 6.**
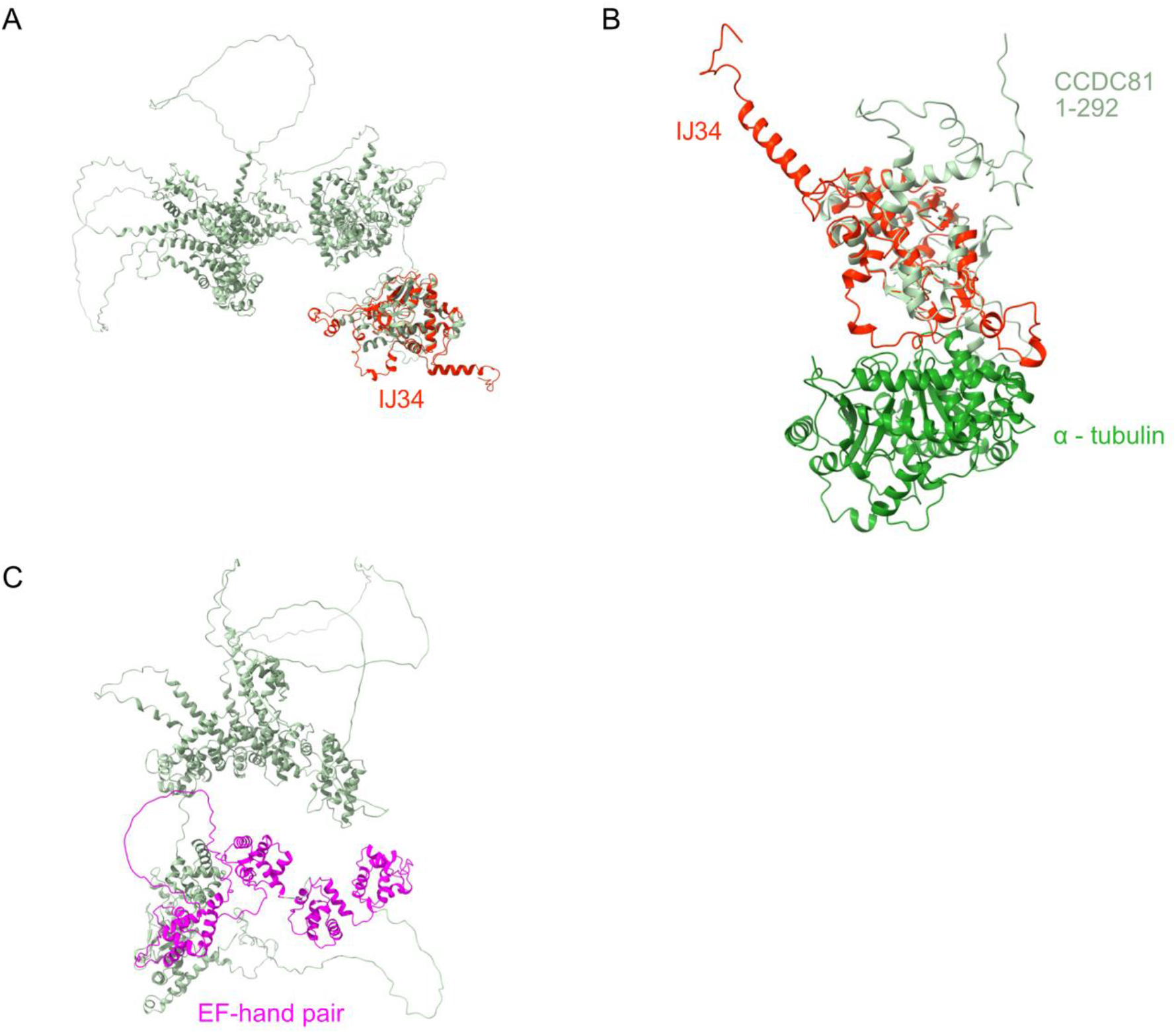
(A) Overlay of the predicted structure of CCDC81 (green) and IJ34 (red). (B) Overlay of the N-terminal domain of CCDC81 shown in (A) and IJ34 along with α-tubulin (green) (C) Predicted structure of CCDC81 with four EF-hand pair domains highlighted in purple.

## Supplementary tables

**Supplementary Table 1.** Data collection parameters for the various datasets used in this study.

| Dataset | *CU428* without membrane | *CU428* with membrane | *CU428* with membrane, lower magnification | *FAP256A/B-KO* |
| --- | --- | --- | --- | --- |
| Magnification | 42,000 | 42,000 | 26,000 | 26,000 |
| Voltage (keV) | 300 | 300 | 300 | 300 |
| Energy filter | Yes | Yes | Yes | Yes |
| Slit width (eV) | 20 | 20 | 20 | 20 |
| Pixel size (Å) | 2.12 | 2.12 | 3.36 | 3.36 |
| Defocus range (µm) | 2 – 4 | 3.5 - 5 | 4.5 - 6 | 3.5 - 5 |
| Tilt range, increment | -60° - 60°, 3° | -60° - 60°, 3° | -60° - 60°, 3° | -60° - 60°, 3° |
| Tilt scheme | Dose symmetric | Dose symmetric | Dose symmetric | Dose symmetric |
| Total dose (e-/Å<sup>2</sup>) | ~164 | ~164 | ~164 | ~164 |
| Number of tilt series used (tip containing) | 136 (66) | 26 (17) | 38 (15) | 33 (29) |

**Supplementary Table 2.** Characteristics of the subtomogram averages shown in this study.

| Structure | Tip CP | Main CP | Tip complex | *FAP256A/B* KO tip CP | A-tubule |
| --- | --- | --- | --- | --- | --- |
| Number of subtomograms | 11,534 | 2,119 | 46 | 4,407 | 3,459 |
| Repeat distance | 8 nm | 32 nm | N/A | 8 nm | 8 nm |
| Resolution (FSC = 0.143) | 8.75 Å | 22.6 Å |  | 23 Å |  |
| Symmetry | C1 | C1 | C1 | C1 | C1 |

**Supplementary Table 3.** List of proteins significantly reduced from FAP256A/B-KO cilia compared to WT and with 4 times fold reduction.

| Uniprot ID | Name | Amino acids | Human homolog? |
| --- | --- | --- | --- |
| Q22Y55 | Phosphatase 2A | 579 | PP2A |
| Q229R5 | Ser/Thr phosphatase | 313 | PP2A |
| I7LX00 | Group 1 truncated haemoglobin | 121 |  |
| I7LVY1 | Kinesin motor catalytic domain protein | 1066 | Kif16B |
| Q22D06 | WD domain, G-beta repeat protein | 2585 | dynein assembly factor with WDR repeat domains |
| Q24CF9 | WD domain, G-beta repeat protein | 2207 | dynein assembly factor with WDR repeat domains |
| I7M9E4 | ATP:guanido phosphotransferase, carboxy-terminal domain protein | 370 | Creatine kinase |
| Q23DC5 | Thioredoxin glutathione reductase | 491 | Thioredoxin reductase |
| I7LU65 | C2 domain protein | 841 |  |
| I7MMW4 | STOP Protein | 592 |  |
| Q22F01 | Phospholipid scramblase | 277 |  |

